# SpaAxFormer: Fine-Grained Token Modeling of Spatial Transcriptomics with Axial Transformers

**DOI:** 10.64898/2026.11.10.765756

**Authors:** Chan Lee, Xiaohui Xie

**Affiliations:** Department of Computer Science, University of California, Irvine, Irvine, CA, USA

## Abstract

Spatial transcriptomics combines gene-expression measurements with spatial relationships between cells. We investigate how spatial context can be incorporated into gene-resolved representations while preserving explicit input-gene correspondence. SpaAxFormer is a spatially constrained factorized formulation that maintains contextual cell–gene states throughout the encoder. It first contextualizes genes within each cell and then exchanges corresponding-gene states across spatial neighbors, while a separate CLS representation provides a cell-level summary. Contextual gene states support gene-expression reconstruction, while CLS-conditioned reconstruction and contrastive learning provide complementary supervision for the cell-level representation.

We evaluate SpaAxFormer on gene-expression imputation and spatial-domain identification across multiple spatial-transcriptomics technologies. Compared with the included baselines, imputation results were most favorable on dorsolateral prefrontal cortex (DLPFC) datasets and more mixed on additional technologies, while post-hoc spatial refinement consistently improved mean ARI on the Visium spatial-domain datasets where it was applied. In a single-slice component-addition analysis, mean clustering ARI increased as CLS-conditioned reconstruction and contrastive supervision were added. Together, these results show that the same gene-resolved formulation can support complementary gene-level prediction and cell-level spatial representation while preserving explicit input-gene correspondence.

## 1 Introduction

Spatial transcriptomics (ST) measures gene expression together with spatial location, enabling molecular variation to be studied in tissue context. Depending on the measurement platform, observations may correspond to individual cells or multicellular spots. This combination of molecular and spatial information creates a central representation-learning challenge: how to model relationships within an expression profile together with relationships among spatially neighboring observations.

A major line of ST methods has focused on incorporating spatial context into cell- or spot-level representations. Probabilistic and graph-based approaches use spatial adjacency, often together with expression similarity, to support spatial-domain identification [10, 8, 15, 12]. Spatial relationships have also been incorporated into gene-expression imputation [9]. These approaches establish the value of neighborhood information for spatial representation learning. In many such formulations, however, the primary unit exchanged across the spatial graph is a cell- or spot-level representation, while gene-level information contributes through the associated expression profile.

Gene-resolved representation learning provides a complementary direction. Gene-token models such as scGPT represent individual genes explicitly and learn contextual representations across an expression profile [6]. Outside spatial transcriptomics, GRN-Transformer organizes single-cell RNA-seq data as cell–gene representations and alternates attention along the gene and cell axes for gene-regulatory-network inference [21]. Its conventional cell-attention module mixes corresponding-gene states across cells using a cell-to-cell attention map shared across genes, so the cross-cell attention weights are computed at the cell-pair level rather than separately for each gene. In nonspatial single-cell transcriptomics, TABULA models expression as a cell-by-gene table, combining gene-level reconstruction with cell-level contrastive learning [7].

Recent ST approaches have increasingly brought gene-level and spatial context together. scGPT-spatial extends a gene-token foundation model to spatial transcriptomics through continual pretraining with spatially aware sampling and a neighborhood-based objective [26]. OrthoST alternates attention across spots and projected expression-feature dimensions within spatially defined local groups, then pools the resulting states into spot-level embeddings [28]. TERRA constructs neighborhood sequences of gene tokens from an index cell and nearby cells, producing spatially contextualized gene-token representations [3]. stFormer incorporates information from the spatial environment through cross-attention with niche-associated gene inputs [4]. Together, these developments establish the integration of gene-level information and spatial context as an emerging direction.

For gene-level prediction, maintaining correspondence between contextual states and retained input genes allows spatial context to be incorporated while retaining gene-specific readouts. This motivates a more specific design question: can within-cell gene context and spatial-neighborhood context be integrated in such a representation while also supporting a cell-level summary?

To investigate this question, SpaAxFormer maintains a contextual state for each cell–gene entry (*i, j*), so that the representation remains associated with gene *j* in cell *i* as information is exchanged along two complementary axes (Fig. 1). Within-cell attention contextualizes gene states within each expression profile, followed by spatially restricted cross-cell attention computed separately for each retained input gene. A separate CLS representation provides a cell-level summary alongside the gene-resolved states. These complementary representation levels support gene-expression prediction and spatial-domain identification.

**Figure 1.**
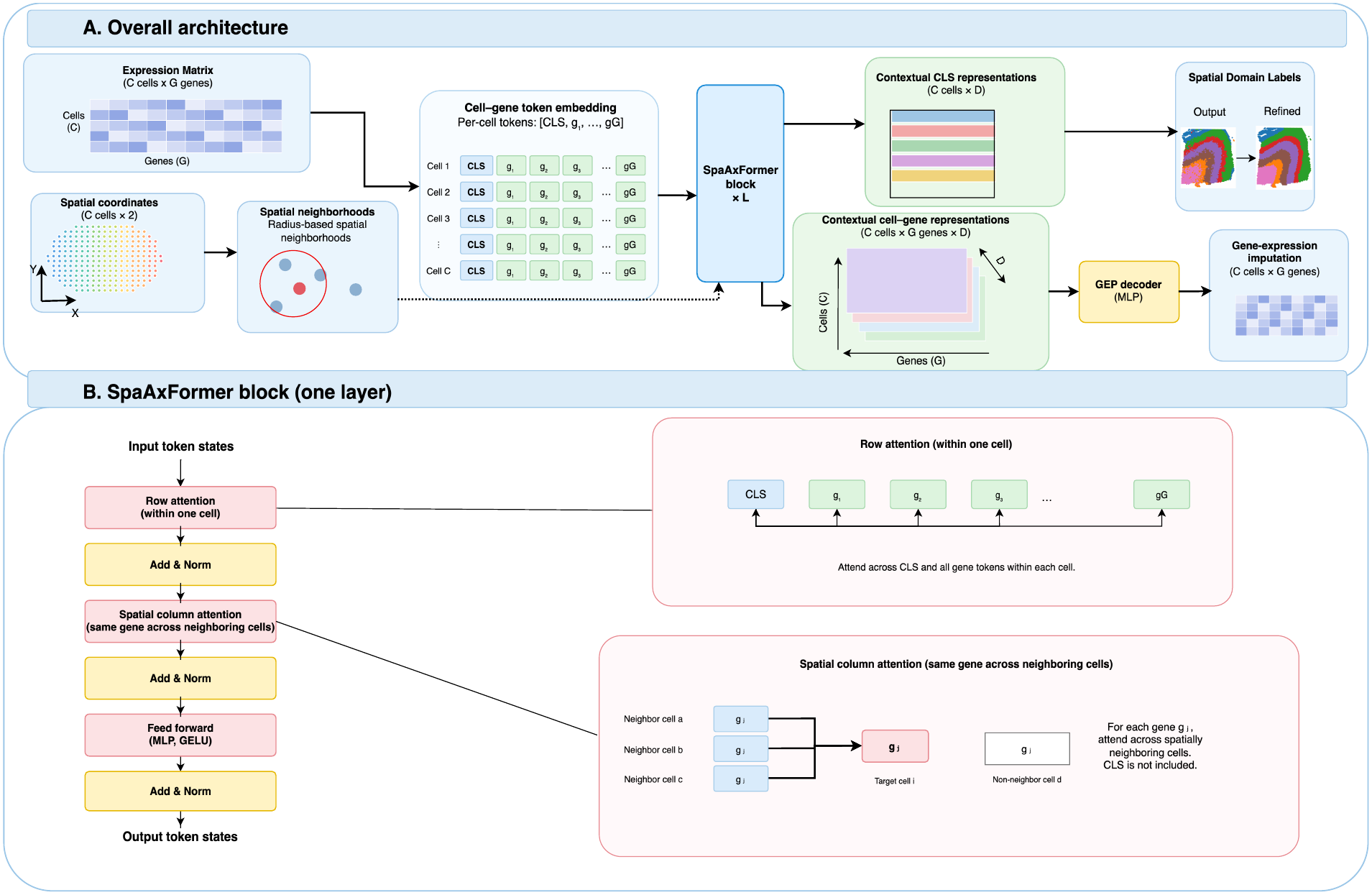
SpaAxFormer architecture and contextual information exchange. (A) Gene-expression inputs form gene-indexed tokens with one CLS position per cell, while spatial coordinates define neighborhoods used by the encoder. Repeated SpaAxFormer blocks produce contextual CLS representations for spatial-domain identification and contextual cell–gene representations for gene-expression imputation through the GEP decoder. (B) Each block applies row attention, spatial column attention, and a feed-forward network with GELU, each followed by residual addition and layer normalization. Row attention includes CLS and all gene tokens within a cell. For each gene, spatial column attention updates the focal cell’s gene representation using corresponding-gene representations from its spatial neighbors; CLS is excluded from this operation.

Training provides supervision at both representation levels. Contextual gene states are used for masked gene-expression reconstruction, while CLS-conditioned reconstruction and contrastive learning provide additional cell-level supervision in spatial-domain configurations. We evaluate SpaAxFormer for gene-expression imputation and spatial-domain identification across multiple spatial-transcriptomics technologies. We further conduct an ablation study on a single dorsolateral prefrontal cortex (DLPFC) slice to examine how additional training objectives affect the learned cell-level representation.

This work makes three contributions:

1. **Gene-resolved and cell-level representations.** SpaAxFormer investigates a gene-resolved spatial formulation that maintains contextual states associated with individual cell–gene entries, together with a separate CLS representation that provides a cell-level summary.
2. **Organization of contextual information exchange.** SpaAxFormer organizes contextual exchange through within-cell attention among gene representations followed by spatially constrained attention among corresponding input-gene representations across neighboring cells, with cross-cell attention weights computed separately for each retained input gene. This organization preserves gene-specific readouts after spatial contextualization while allowing cell-level information to be summarized separately through CLS.
3. **Empirical evaluation.** We evaluate this formulation for complementary gene-level and cell-level tasks, including gene-expression imputation and spatial-domain identification, across multiple spatial-transcriptomics technologies. We further examine the contribution of additional training objectives through a controlled single-slice ablation.

## 2 Methods

### 2.1 Problem formulation

Let *X* ∈ ℝ^*C×G*^ denote the preprocessed expression matrix supplied to the model, with *C* spatial observations and *G* retained input genes, and let *P* ∈ ℝ^*C×*2^ contain their spatial coordinates. Throughout Methods, we use ‘cell’ to denote a spatial observation, which may correspond to an individual cell or a multicellular spot depending on the measurement technology.

Given (*X, P*), SpaAxFormer learns an encoder

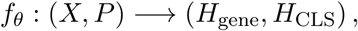

with

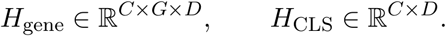

Here, *D* is the embedding dimension. We denote the contextual representation of input gene *j* in cell *i* by **h**_*ij*_ ∈ ℝ^*D*^, and the contextual CLS representation of cell *i* by **h**_CLS,*i*_ ∈ ℝ^*D*^.

These two representation levels support complementary downstream tasks. Contextual gene representations are used to predict held-out expression values for selected cell–gene entries, whereas the CLS representations provide the cell-level embeddings used for spatial-domain identification. Spatial coordinates determine which cells may exchange information during spatial contextualization; the coordinates themselves are not embedded directly as token features.

The following sections describe how cell–gene input representations are constructed, how information is exchanged within cells and across spatial neighbors, and how the resulting representations are supervised (Fig. 1).

### 2.2 Cell–gene input representations

Before contextualization, SpaAxFormer represents each entry of the expression matrix as a token associated with one particular gene in one particular cell. Thus, token (*i, j*) remains tied to cell *i* and input gene *j* throughout the encoder. Its initial representation combines information about which cell and gene it represents with information about the measured expression value.

For cell *i* and gene *j*, the input token **x**_*ij*_ ∈ ℝ^*D*^ contains three components. Learned cell-identity and gene-identity embeddings provide the identity representation

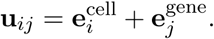

Expression is represented in two complementary forms to provide both continuous expression magnitude and a discretized expression-level representation. A continuous magnitude representation is obtained by applying a two-layer multilayer perceptron *f*_mag_ to the per-gene expression z-score

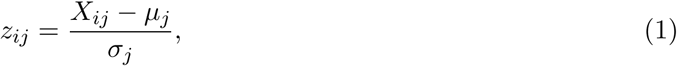

where *µ*_*j*_ and *σ*_*j*_ are the per-gene mean and population standard deviation fitted from the preprocessed expression matrix. In parallel, a learned bin embedding 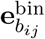 represents a quantile-derived expression level for the same preprocessed value *X*_*ij*_. Expression-bin boundaries are fitted from the nonzero preprocessed expression values *X*_*ij*_ outside *ℋ*.

With *Q* expression bins, observed zeros are assigned to bin 0, while observed nonzero values are assigned to bins 1, …, *Q* − 1. Masking removes expression-dependent information from a token while preserving its cell and gene identity. This distinction is important because measured zero expression is a valid input and must remain distinguishable from an entry hidden from the model. An unmasked token therefore contains its identity, bin, and magnitude components, whereas a masked token uses a distinct learned mask-bin embedding and has no magnitude contribution:

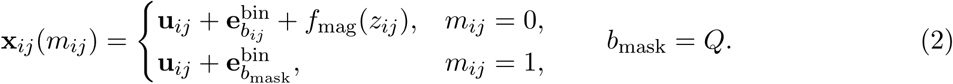

Here, *m*_*ij*_ = 1 denotes an expression entry hidden from the model input. For masked entries, the magnitude contribution is explicitly set to zero rather than obtained by evaluating *f*_mag_ at a zero input. Section 2.4 specifies how reconstruction-training masks and any fixed held-out entries determine which inputs are hidden.

Each cell’s gene tokens are accompanied by the same learned initial CLS vector **h**_CLS_ ∈ ℝ^*D*^:

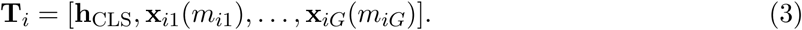

For *K* cells included in an encoder batch, these sequences form a tensor of shape [*K, G* + 1, *D*]. Position 0 contains CLS, while positions 1, …, *G* retain correspondence to the input genes throughout the encoder. The initial CLS vector is shared across cells; after contextualization, its state becomes cell-specific.

### 2.3 Factorized contextual information exchange

The encoder applies *L* SpaAxFormer blocks to states represented by *H* ∈ ℝ^*K×*(*G*+1)*×D*^, where *K* is the number of included cells, each with one CLS position and *G* input-gene positions. Each block factorizes contextual exchange along two axes: row attention holds the cell fixed and exchanges information across CLS and gene states; spatial column attention holds the input-gene identity fixed and exchanges information across spatially permitted cells. This organization combines within-cell and spatial context while preserving cell–gene correspondence. Similar factorized contextualization along cell and gene axes has also been used in GRN-Transformer [21].

Within each block, row attention, spatial column attention, and the feed-forward transformation each use residual addition followed by layer normalization (post-norm) [25]. We write 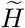 for the states after row attention, residual addition, and layer normalization. A position-wise feed-forward network follows the attention sublayers.

#### 2.3.1 Within-cell row attention

For cell *i*, row attention operates on *H*_*i*_ ∈ ℝ^(*G*+1)*×D*^, containing CLS and all gene states. Each gene state can incorporate information from other genes in the same cell, while CLS can incorporate information across the cell’s gene representations. For one head of dimension *d*_*h*_, with projection matrices in 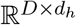,

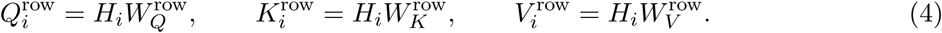

The attention weights and weighted output are

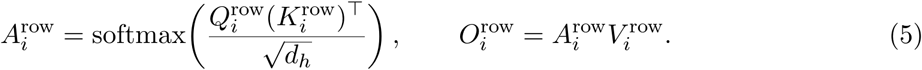

Softmax acts across the *G* + 1 key positions, and attention is computed separately for each cell.

#### 2.3.2 Corresponding-gene spatial attention

After row attention, let 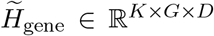 denote the restriction of 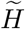 to its *G* gene positions, excluding the CLS position. For target cell *i* and input gene *j*, spatial attention updates the gene-*j* representation using the corresponding gene-*j* representations from spatially permitted cells.

Spatial coordinates define a radius-based graph, with a nearest non-self neighbor added when no other cell lies within the radius and self-loops included for spatial attention. Let *A*_*i*_ = *N*_*r*_(*i*) ∪ *{i}* denote the permitted cells for target cell *i*, including the target itself, where *N*_*r*_(*i*) is the non-self spatial neighborhood. We denote the target state by 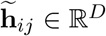 and collect the corresponding gene-*j* states from the permitted cells in 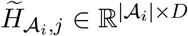.

For one attention head of dimension *d*_*h*_, with projection matrices 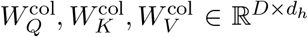, the query is formed from the target cell–gene state, while keys and values are formed from the corresponding gene states of the permitted cells:

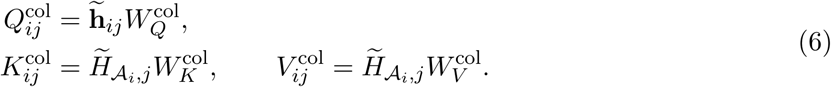

The attention weights and weighted output are

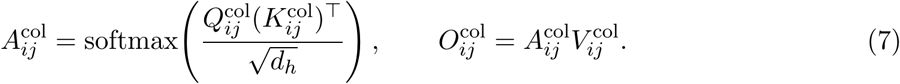

Softmax is applied across the permitted cells in *A*_*i*_. Only the target cell’s gene-*j* state receives the spatial update. The operation is applied separately for each input gene, so spatial exchange preserves input-gene correspondence.

The row-attention output is added to its input and layer-normalized before spatial column attention. The spatial-attention output likewise uses residual addition followed by layer normalization. CLS is then reattached to the updated gene states, and a position-wise feed-forward sublayer with residual addition and layer normalization completes the block. Spatially updated gene states can influence CLS through row attention in subsequent blocks.

After the final encoder block, each contextual gene state **h**_*ij*_ remains associated with input gene *j* in cell *i*, while **h**_CLS,*i*_ provides the cell-level state. These contextual representations are used by the learning objectives described below.

### 2.4 Training procedure and learning objectives

Training supervises the gene-resolved and cell-level representations through complementary objectives. Gene-expression prediction (GEP) reconstructs masked expression values directly from contextual gene states. When enabled, CLS-conditioned gene-expression prediction (GEPC) reconstructs the same targets from the contextual CLS representation together with the corresponding cell– gene identity. When enabled, contrastive learning provides additional supervision for the CLS representations using spatial and expression relationships.

#### 2.4.1 Masking and neighborhood-expanded training

Training combines reconstruction masking with neighborhood-expanded batches. For reconstruction, selected expression entries are hidden from the encoder and retained as prediction targets. If a fixed evaluation holdout *ℋ* is used, those entries remain hidden throughout training and are reserved for evaluation; otherwise, *ℋ* = ∅.

Per-gene z-score statistics are fitted across cells in the input dataset, including zero expression values but excluding entries in *ℋ*. Expression-bin boundaries are fitted from the nonzero expression values outside *ℋ*.

At epoch *t*, a separate reconstruction set *ℛ*^(*t*)^ is sampled from the remaining eligible entries. The entries hidden from the encoder are therefore

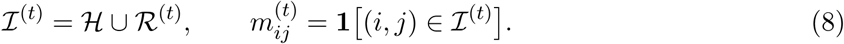

Spatial attention additionally requires neighboring cells to be available when a target cell is updated. At each epoch, cells are randomly permuted and partitioned into target sets *T*_*b*_ of at most *B*_target_ cells, so that each cell appears once as a target. Each target set is expanded with its graph-defined non-self spatial neighbors:

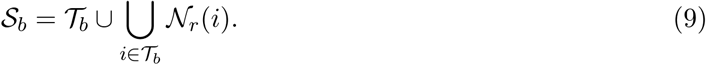

Cells in *S*_*b*_ *\ T*_*b*_ provide spatial context for the targets.

Within an expanded batch, row attention contextualizes the gene and CLS states of all included cells. The resulting states of neighboring cells serve as context for spatial column attention, while spatial-column and feed-forward updates are applied to the target cells.

Reconstruction supervision is restricted to sampled training entries in the target cells:

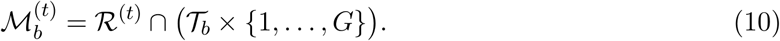

The entries in 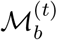 provide the shared reconstruction targets for GEP and GEPC.

#### 2.4.2 Gene-expression prediction

GEP and GEPC predict the masked preprocessed expression values *X*_*ij*_ through different representation levels. GEP predicts each target from its contextual gene state **h**_*ij*_, whereas GEPC requires the cell-level CLS representation **h**_CLS,*i*_ to support gene-specific expression prediction.

For GEP, a prediction head maps each contextual gene state independently to a scalar expression estimate, 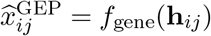. Writing 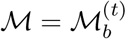 for the current batch’s reconstruction target set, the GEP loss is

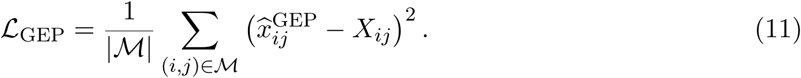

Motivated by scGPT-spatial’s neighborhood-conditioned expression prediction [26], GEPC follows the cell-embedding/gene-query decoding principle of scGPT [6], using the CLS representation of the same cell together with a cell–gene identity query. For GEPC, the cell–gene identity representation **u**_*ij*_ specifies which gene is queried from the cell-level CLS state. Here, 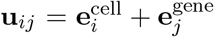 reuses the tokenization identity embeddings without the bin or magnitude components. The identity representation is transformed into a query vector,

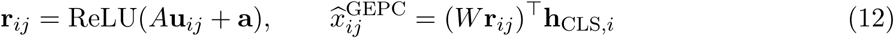

The transformed identity query therefore provides a gene-specific decoding vector whose dot product with the CLS representation of the same cell yields the expression prediction, with *A, W*, and **a** shared across cells and genes. Here, *A, W* ∈ ℝ^*D×D*^ are learned matrices and **a** ∈ ℝ^*D*^ is a learned bias. GEPC uses the same masked target set:

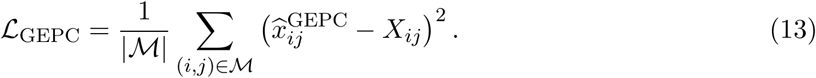

#### 2.4.3 PCA-based weighted multi-positive contrastive learning

GEP and GEPC supervise expression reconstruction, while the contrastive objective directly organizes the cell-level CLS representations using spatial and expression information. Following SpaGIC’s neighborhood-based contrastive formulation [14], spatial neighbors define the candidate positive set for each anchor. Within this set, expression-PCA similarity weights each neighbor’s contribution, allowing spatially adjacent cells with dissimilar expression profiles to exert weaker positive influence. The optimized similarities themselves are computed between the learned CLS representations.

For a target cell *i* ∈ *T*_*b*_ serving as an anchor, graph neighbors present in the expanded batch form the positive set:

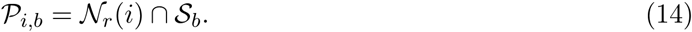

All non-self cells in *S*_*b*_ form the comparison pool in the contrastive denominator; *P*_*i,b*_ identifies the subset treated as positives.

Let **p**_*i*_ denote the *d*_expr_-dimensional expression-PCA representation of cell *i*, derived from the cell-by-gene expression matrix. For an anchor *i* and comparison cell *j*, learned CLS similarity and expression-PCA similarity are

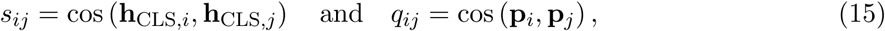

Each positive receives weight *w*_*ij*_ = sigmoid(*αq*_*ij*_), where *α* controls positive-weight sharpness. These weights modulate positive contributions in the numerator of the multi-positive InfoNCE loss [23]. The denominator contains all non-self comparison cells, including positives, with similarities scaled by temperature *τ* :

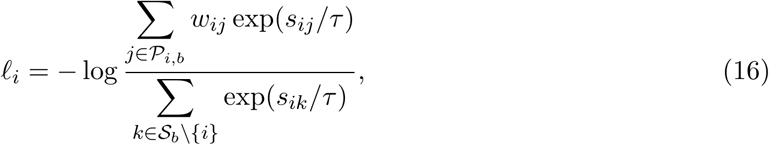

The batch loss averages over target anchors with at least one positive, *V*_*b*_ = *{i* ∈ *T*_*b*_ : |*P*_*i,b*_| *>* 0*}*:

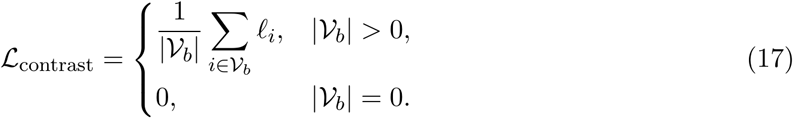

##### Combined training objective

The general training objective combines gene-level reconstruction, CLS-conditioned reconstruction, and contrastive supervision, with component weights *λ*_GEP_, *λ*_GEPC_, and *λ*_contrast_:

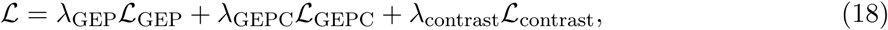

### 2.5 Gene-expression imputation

For gene-expression imputation, a fixed holdout set *ℋ* of measured cell–gene entries is hidden from the encoder throughout training and reserved for evaluation. SpaAxFormer uses the GEP-only objective,

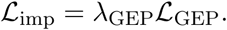

The GEP decoder predicts each held-out value from its contextual gene representation, producing 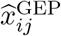 for (*i, j*) ∈ *ℋ*.

The predicted values are evaluated against the corresponding held-out measurements *X*_*ij*_. Task-specific preprocessing, masking, and evaluation protocols are described in Section 3.2.

### 2.6 Spatial-domain identification

After representation learning, the contextual CLS states form a cell-level embedding matrix. These embeddings are reduced by PCA and clustered with a Gaussian mixture model using *mclust* [20] to obtain initial spatial-domain labels. This PCA is applied to the learned CLS representations and is distinct from the expression PCA used to weight contrastive positives.

For spatial-domain identification, SpaAxFormer is trained with the full objective combining GEP, GEPC, and PCA-based weighted multi-positive contrastive learning (Eq. 18). Task-specific clustering settings and the component-addition ablation protocol are described in Sections 3.3 and 3.4, respectively.

#### 2.6.1 Post-hoc spatial label refinement

Following the spatial refinement strategy used in GraphST [15], we optionally refine the domain assignments obtained from *mclust* using local spatial context. For each cell, the 50 nearest spatial neighbors are identified by Euclidean coordinate distance, and the cell is assigned the most frequent cluster label among these neighbors. Refinement is applied once after clustering and does not alter the learned CLS representations or model parameters.

## 3 Experimental Design

### 3.1 Datasets and evaluation overview

SpaAxFormer is evaluated on two downstream tasks: gene-expression imputation and spatial-domain identification. Imputation uses six DLPFC slices (151507, 151508, 151509, 151510, 151669, and 151670) [17, 18], Stereo-seq [5], and Slide-seq V2 [22]. Spatial-domain identification uses eight DLPFC slices (151507, 151508, 151509, 151510, 151673, 151674, 151675, and 151676), BC1 [1], and STARMap [29]. The evaluated datasets span multiple sequencing- and imaging-based spatial-transcriptomics technologies, tissues, and species. Task-specific preprocessing, masking, training, and evaluation settings are described in the following subsections.

Across the selected SpaAxFormer runs, training used AdamW with a learning rate of 10^−3^, weight decay 5 *×* 10^−4^, four encoder layers, one configured attention head, a maximum of 15 target cells per neighborhood-expanded batch, and no learning-rate scheduler. Imputation models were trained for 20 epochs; DLPFC, BC1, and STARMap clustering models were trained for 150, 50, and 400 epochs, respectively.

Baseline values are reproduced from the corresponding published benchmark studies rather than obtained by rerunning all comparison methods under a common preprocessing and training pipeline; comparisons therefore retain the protocols of their respective source studies.

### 3.2 Gene-expression imputation evaluation

Gene-expression imputation uses the Impeller-preprocessed expression matrices and fixed test masks [9], without repeating normalization or highly-variable-gene selection. The fixed test entries are withheld from model inputs and reconstruction supervision and are excluded when fitting pergene z-score statistics and expression-bin boundaries. SpaAxFormer uses the GEP-only configuration with CLS retained, while GEPC and contrastive learning are disabled. During training, 30% of eligible observed nonzero entries outside the fixed test holdout are masked for reconstruction.

Performance is evaluated on the fixed held-out entries using L1 distance, root mean squared error (RMSE), and cosine similarity. SpaAxFormer results are reported as mean *±* sample standard deviation (*ddof=1*) across ten fixed test masks. Comparisons include MAGIC [24], scVI [16], scGNN [27], Tangram [2], STLearn [19], STAGATE, and Impeller, using the published Impeller benchmark values [9].

### 3.3 Spatial-domain identification evaluation

Spatial-domain identification is evaluated on DLPFC slices, BC1, and STARMap. For DLPFC, preprocessing follows the ADEPT-derived pipeline used for the selected runs [12]: genes are filtered until the nonzero expression fraction exceeds 0.14, 3,000 highly variable genes are requested using *seurat v3*, counts are normalized to 10^4^ per cell and log-transformed, and the selected genes are retained.

Training uses no fixed expression holdout and masks 30% of expression entries for reconstruction. SpaAxFormer uses the full GEP + GEPC + PCA-based weighted multi-positive InfoNCE objective. The contrastive weight is 0.02 for DLPFC and BC1 and 2.5 for STARMap. Spatial radii are 150 for DLPFC and BC1 and 450 for STARMap. Expression PCA retained 100 principal components for computing contrastive positive weights in all DLPFC, BC1, and STARMap clustering experiments.

After training, CLS representations were reduced to min(30, *D, C* − 1) principal components and clustered with *mclust* (Section 2.6). Post-hoc spatial refinement is applied to the Visium spatial-domain datasets (DLPFC and BC1) without retraining the encoder. Given its single-cell spatial resolution, STARMap is evaluated using the unrefined clustering assignments. Performance is measured by adjusted Rand index (ARI) against the supplied domain annotations. SpaAxFormer results are reported as mean *±* sample standard deviation (*ddof=1*) across five random seeds. Comparisons include ADEPT, STAGATE, CCST [13], SEDR [30], SpaGCN, and BayesSpace [31], using the published ADEPT comparison values [12], which were reported over 20 runs under its evaluation protocol.

### 3.4 Clustering ablation

We evaluate the contribution of additional training objectives on DLPFC slice 151675 using three configurations: GEP, GEP + GEPC, and the full objective with contrastive learning. All configurations use the same encoder architecture, experimental setup, and five random seeds, with CLS retained as the clustering representation. The reported ablation uses unrefined ARI to compare the training objectives before post-hoc spatial label refinement.

| Configuration | Training objective |
|---|---|
| GEP | 5ℒ<sub>GEP</sub> |
| GEP + GEPC | 5ℒ<sub>GEP</sub> + 5ℒ<sub>GEPC</sub> |
| Full objective | 5ℒ<sub>GEP</sub> + 5ℒ<sub>GEPC</sub> + 0.02ℒ<sub>contrast</sub> |

## 4 Results

### 4.1 Gene-expression imputation across DLPFC and additional ST technologies

To evaluate whether the contextual gene states learned by SpaAxFormer retain information useful for gene-expression reconstruction, we measured imputation accuracy on held-out expression entries across six DLPFC slices and two additional spatial-transcriptomics datasets, Stereo-seq and Slide-seq V2. For this task, SpaAxFormer used GEP to predict masked expression values from the contextual cell–gene representations.

Across the six DLPFC slices, the learned gene-level representations consistently supported reconstruction of held-out expression. SpaAxFormer had lower reported mean L1 distance than Impeller [9] on all six slices, with closely matched RMSE and cosine similarity (Tables 2, 4, and 3). For example, on slice 151510, reported mean L1 was 0.239 for SpaAxFormer compared with 0.254 for Impeller. Across DLPFC, SpaAxFormer also had lower reported mean RMSE on four slices and tied on two, while cosine similarity was higher on three slices and tied on the remaining three at the reported precision.

**Table 1:**
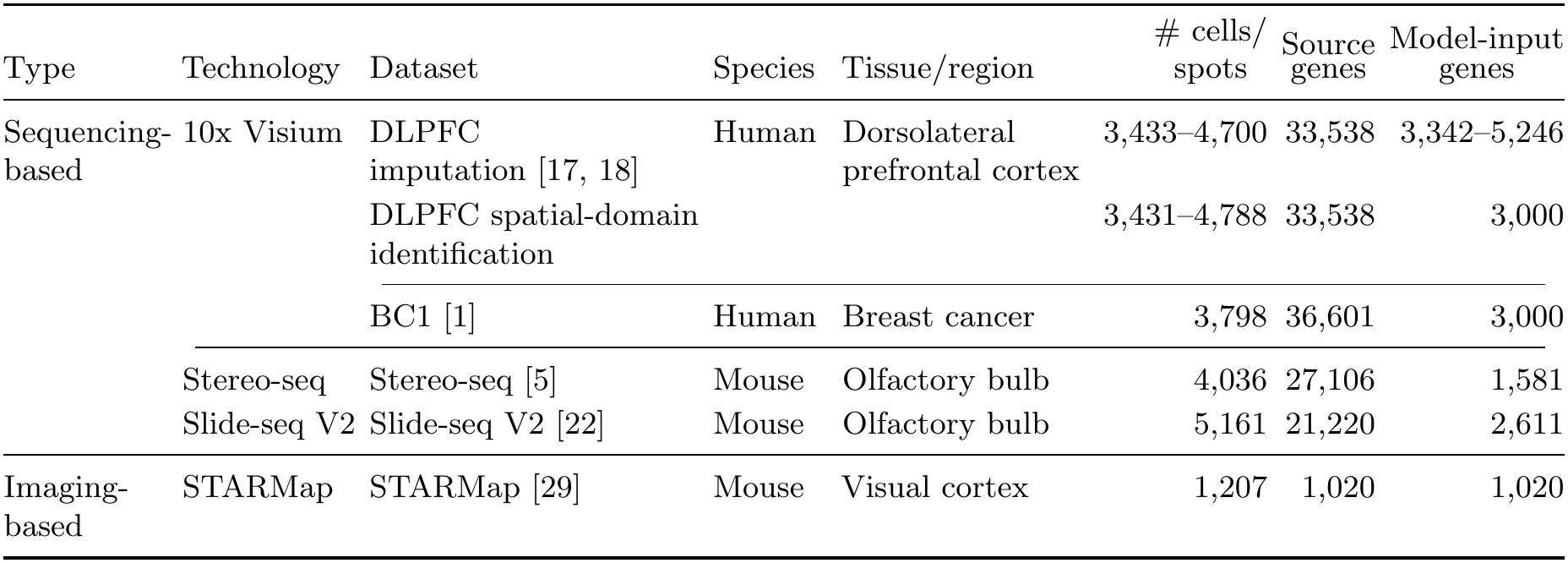
Summary of spatial transcriptomics datasets evaluated in this study. Cell/spot counts describe the evaluated inputs. Source genes denote the number of features in the source expression matrices before gene reduction, and model-input genes denote the genes retained in the matrices supplied to SpaAxFormer. Ranges summarize variation across slices.

**Table 2:** L1 distance (↓) for gene-expression imputation. Mean *±* SD across 10 fixed test masks. SpaAxFormer uses GEP; baseline values are from the Impeller comparison. *NA* reproduces unavailable source entries.

| Method | 151507 | 151508 | 151509 | 151510 | 151669 | 151670 |
|---|---|---|---|---|---|---|
| MAGIC | 0.779±0.001 | 0.825±0.001 | 0.787±0.000 | 0.664±0.001 | 0.795±0.001 | 0.692±0.000 |
| scVI | 0.794±0.004 | 0.838±0.006 | 0.800±0.002 | 0.670±0.003 | 0.810±0.003 | 0.696±0.005 |
| scGNN | 0.583±0.011 | 0.665±0.085 | 0.589±0.011 | 0.584±0.004 | 0.550±0.006 | 0.532±0.009 |
| Tangram | 1.691±0.001 | 1.811±0.001 | 1.689±0.000 | 1.420±0.000 | 1.728±0.001 | 1.474±0.000 |
| STLearn | 1.333±0.001 | 1.423±0.001 | 1.332±0.001 | 1.148±0.001 | 1.369±0.002 | 1.206±0.001 |
| STAGATE | 0.297±0.001 | 0.300±0.002 | 0.295±0.005 | 0.294±0.004 | 0.274±0.005 | 0.278±0.002 |
| Impeller | 0.248±0.001 | 0.252±0.003 | 0.247±0.003 | 0.254±0.003 | 0.242±0.004 | 0.237±0.001 |
| SpaAxFormer GEP | 0.241±0.007 | 0.243±0.0097 | 0.237±0.0089 | 0.239±0.0102 | 0.229±0.0087 | 0.234±0.006 |
| Method | Stereo-seq | Slide-seq V2 |  |  |  |  |
| MAGIC | 1.324±0.001 | 1.080±0.000 |  |  |  |  |
| scVI | 1.442±0.005 | 1.127±0.006 |  |  |  |  |
| scGNN | 0.819±0.240 | 0.664±0.018 |  |  |  |  |
| Tangram | 2.899±0.001 | 2.185±0.000 |  |  |  |  |
| STLearn | NA | NA |  |  |  |  |
| STAGATE | 0.289±0.006 | 0.502±0.007 |  |  |  |  |
| Impeller | 0.190±0.005 | 0.292±0.005 |  |  |  |  |
| SpaAxFormer GEP | 0.272±0.0434 | 0.297±0.044 |  |  |  |  |

**Table 3:**
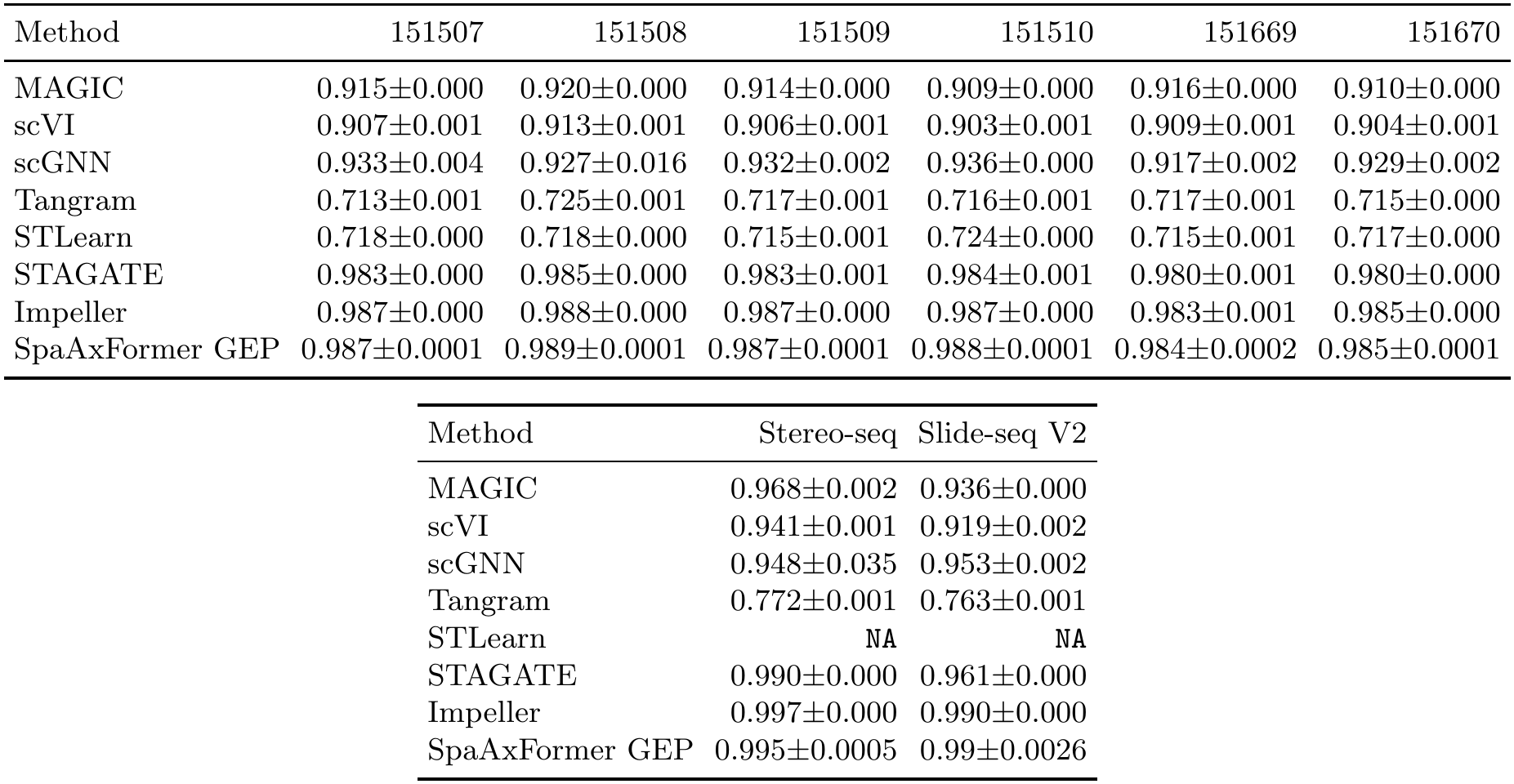
Cosine similarity (↑) for gene-expression imputation. Mean *±* SD across 10 fixed test masks. SpaAxFormer uses GEP; baseline values are from the Impeller comparison. *NA* reproduces unavailable source entries.

| Method | 151507 | 151508 | 151509 | 151510 | 151669 | 151670 |
|---|---|---|---|---|---|---|
| MAGIC | 0.915±0.000 | 0.920±0.000 | 0.914±0.000 | 0.909±0.000 | 0.916±0.000 | 0.910±0.000 |
| scVI | 0.907±0.001 | 0.913±0.001 | 0.906±0.001 | 0.903±0.001 | 0.909±0.001 | 0.904±0.001 |
| scGNN | 0.933±0.004 | 0.927±0.016 | 0.932±0.002 | 0.936±0.000 | 0.917±0.002 | 0.929±0.002 |
| Tangram | 0.713±0.001 | 0.725±0.001 | 0.717±0.001 | 0.716±0.001 | 0.717±0.001 | 0.715±0.000 |
| STLearn | 0.718±0.000 | 0.718±0.000 | 0.715±0.001 | 0.724±0.000 | 0.715±0.001 | 0.717±0.000 |
| STAGATE | 0.983±0.000 | 0.985±0.000 | 0.983±0.001 | 0.984±0.001 | 0.980±0.001 | 0.980±0.000 |
| Impeller | 0.987±0.000 | 0.988±0.000 | 0.987±0.000 | 0.987±0.000 | 0.983±0.001 | 0.985±0.000 |
| SpaAxFormer GEP | 0.987±0.0001 | 0.989±0.0001 | 0.987±0.0001 | 0.988±0.0001 | 0.984±0.0002 | 0.985±0.0001 |
| Method | Stereo-seq | Slide-seq V2 |  |  |  |  |
| MAGIC | 0.968±0.002 | 0.936±0.000 |  |  |  |  |
| scVI | 0.941±0.001 | 0.919±0.002 |  |  |  |  |
| scGNN | 0.948±0.035 | 0.953±0.002 |  |  |  |  |
| Tangram | 0.772±0.001 | 0.763±0.001 |  |  |  |  |
| STLearn | NA | NA |  |  |  |  |
| STAGATE | 0.990±0.000 | 0.961±0.000 |  |  |  |  |
| Impeller | 0.997±0.000 | 0.990±0.000 |  |  |  |  |
| SpaAxFormer GEP | 0.995±0.0005 | 0.99±0.0026 |  |  |  |  |

**Table 4:** RMSE (↓) for gene-expression imputation. Mean *±* SD across 10 fixed test masks. SpaAxFormer uses GEP; baseline values are from the Impeller comparison. *NA* reproduces unavailable source entries.

| Method | 151507 | 151508 | 151509 | 151510 | 151669 | 151670 |
|---|---|---|---|---|---|---|
| MAGIC | 0.917±0.001 | 0.972±0.001 | 0.929±0.000 | 0.792±0.000 | 0.936±0.001 | 0.824±0.000 |
| scVI | 0.940±0.005 | 0.993±0.006 | 0.949±0.003 | 0.803±0.003 | 0.959±0.003 | 0.834±0.005 |
| scGNN | 0.755±0.016 | 0.850±0.096 | 0.762±0.011 | 0.755±0.002 | 0.717±0.007 | 0.686±0.010 |
| Tangram | 1.768±0.001 | 1.889±0.001 | 1.767±0.000 | 1.503±0.000 | 1.804±0.001 | 1.557±0.000 |
| STLearn | 1.516±0.001 | 1.629±0.001 | 1.521±0.001 | 1.300±0.001 | 1.556±0.002 | 1.362±0.001 |
| STAGATE | 0.384±0.002 | 0.393±0.002 | 0.379±0.007 | 0.380±0.007 | 0.357±0.007 | 0.365±0.004 |
| Impeller | 0.337±0.001 | 0.341±0.000 | 0.336±0.000 | 0.340±0.004 | 0.327±0.007 | 0.323±0.000 |
| SpaAxFormer GEP | 0.336±0.0011 | 0.341±0.002 | 0.335±0.0028 | 0.337±0.0033 | 0.321±0.0024 | 0.323±0.0015 |
| Method | Stereo-seq | Slide-seq V2 |  |  |  |  |
| MAGIC | 1.453±0.001 | 1.238±0.001 |  |  |  |  |
| scVI | 1.628±0.005 | 1.307±0.007 |  |  |  |  |
| scGNN | 1.051±0.307 | 0.842±0.021 |  |  |  |  |
| Tangram | 2.970±0.001 | 2.284±0.000 |  |  |  |  |
| STLearn | NA | NA |  |  |  |  |
| STAGATE | 0.485±0.008 | 0.765±0.005 |  |  |  |  |
| Impeller | 0.277±0.002 | 0.391±0.006 |  |  |  |  |
| SpaAxFormer GEP | 0.364±0.0394 | 0.401±0.0482 |  |  |  |  |

Reconstruction behavior differed on the additional spatial-transcriptomics datasets. On Stereo-seq and Slide-seq V2, Impeller had lower reported mean L1 and RMSE, although the differences were small on Slide-seq V2 (0.297 versus 0.292 for L1 and 0.401 versus 0.391 for RMSE). Cosine similarity on Slide-seq V2 was tied at 0.990. At the same time, SpaAxFormer had lower reported mean L1 and RMSE and higher reported mean cosine similarity than STAGATE on both datasets. Together, these results show that the contextual gene states supported competitive reconstruction across the evaluated datasets, with the strongest relative performance on DLPFC and greater variation across the additional technologies.

### 4.2 Spatial-domain identification across DLPFC, BC1, and STARMap

The full-objective SpaAxFormer configuration produces one contextual CLS representation per spatial observation. We clustered these representations after PCA to evaluate whether they organize spatial observations in agreement with annotated tissue domains (Table 5). The evaluation spans eight DLPFC slices and BC1, measured at Visium spot resolution, together with single-cell-resolution STARMap data.

**Table 5:** Spatial-domain identification measured by ARI (↑). SpaAxFormer rows report mean *±* SD across five random seeds. Full denotes GEP + GEPC + PCA-based weighted multi-positive InfoNCE; refined denotes post-hoc spatial label refinement, applied only to the Visium datasets (DLPFC and BC1). The dash for STARMap indicates that refinement was not used in the primary comparison. Baseline values come from the ADEPT comparison and follow its reported 20-run protocol.

| Method | 151507 | 151508 | 151509 | 151510 | 151673 |
|---|---|---|---|---|---|
| SpaAxFormer full | 0.478±0.0229 | 0.348±0.0339 | 0.48±0.022 | 0.428±0.055 | 0.501±0.0566 |
| SpaAxFormer refined | 0.525±0.0346 | 0.407±0.0398 | 0.516±0.045 | 0.498±0.0712 | 0.513±0.062 |
| ADEPT | 0.543 | 0.492 | 0.537 | 0.534 | 0.603 |
| STAGATE | 0.572 | 0.479 | 0.473 | 0.489 | 0.537 |
| CCST | 0.444 | 0.400 | 0.434 | 0.390 | 0.455 |
| SEDR | 0.434 | 0.352 | 0.376 | 0.356 | 0.485 |
| SpaGCN | 0.461 | 0.382 | 0.455 | 0.446 | 0.465 |
| BayesSpace | 0.380 | 0.338 | 0.323 | 0.291 | 0.460 |
| Method | 151674 | 151675 | 151676 | BC1 | STARMap |
| SpaAxFormer full | 0.473±0.0631 | 0.511±0.0519 | 0.393±0.0779 | 0.434±0.0452 | 0.418±0.080 |
| SpaAxFormer refined | 0.491±0.0663 | 0.525±0.0483 | 0.42±0.1033 | 0.513±0.0579 | – |
| ADEPT | 0.510 | 0.649 | 0.586 | 0.563 | 0.611 |
| STAGATE | 0.486 | 0.589 | 0.562 | 0.481 | 0.496 |
| CCST | 0.385 | 0.492 | 0.505 | 0.549 | 0.147 |
| SEDR | 0.452 | 0.451 | 0.474 | 0.507 | 0.230 |
| SpaGCN | 0.403 | 0.400 | 0.340 | 0.572 | 0.379 |
| BayesSpace | 0.320 | 0.352 | 0.335 | 0.489 | 0.262 |

Across the evaluated datasets, the learned CLS representations supported recovery of annotated spatial-domain structure, although agreement varied across settings. Before post-hoc refinement, reported mean ARI ranged from 0.348 to 0.511 across the DLPFC slices and was 0.434 on BC1 and 0.418 on STARMap. Unrefined SpaAxFormer had higher reported mean ARI than CCST, SEDR, and SpaGCN in seven settings each and higher reported mean ARI than BayesSpace in nine. ADEPT and STAGATE provided stronger comparisons: ADEPT had higher reported mean ARI in all ten settings, while SpaAxFormer had higher reported mean ARI than STAGATE on DLPFC slice 151509.

Post-hoc spatial label refinement further increased reported mean ARI in all nine Visium evaluations on which it was applied, including an increase from 0.434 to 0.513 on BC1. After refinement, SpaAxFormer had higher reported mean ARI than STAGATE on DLPFC slices 151509, 151510, and 151674, as well as BC1, while ADEPT remained higher in all nine refined Visium comparisons. STARMap remained evaluated using unrefined assignments, with reported mean ARI of 0.418 compared with 0.496 for STAGATE. Because refinement changes the initial clustering labels without retraining the encoder, these gains reflect a downstream label operation rather than an improvement in the learned CLS representations themselves.

### 4.3 Clustering ablation

To examine how the training objectives relate to the cell-level representation used for spatial-domain identification, we compared three objective configurations on DLPFC slice 151675 while retaining the same encoder architecture and CLS clustering representation. The comparison progressively adds GEPC and then PCA-based weighted multi-positive InfoNCE to the GEP objective.

Under this setup, the resulting CLS-based clustering showed a stepwise increase in mean ARI. Mean ARI was 0.103 with GEP alone, increased to 0.400 with GEP + GEPC, and reached 0.511 with the full objective (Table 6). All three configurations used the same five random seeds and experimental setup, with CLS retained as the clustering representation throughout. On this single DLPFC slice, progressively adding GEPC and contrastive supervision was therefore associated with higher mean ARI for the CLS-based spatial-domain readout.

**Table 6:** Clustering ablation on DLPFC 151675. Values report mean unrefined ARI *±* SD across the same five random seeds. CLS is retained throughout; full denotes GEP + GEPC + PCA-based weighted multi-positive InfoNCE.

| Configuration | ARI ↑ |
|---|---|
| GEP | 0.103±0.0196 |
| GEP + GEPC | 0.4±0.0637 |
| Full objective | 0.511±0.0519 |

## 5 Discussion and Conclusion

SpaAxFormer investigates a gene-resolved approach to spatial representation learning in which contextual states remain associated with individual cell–gene entries while a separate CLS representation summarizes each spatial observation. The experiments show that these two representation levels can support complementary tasks within the same formulation: contextual gene states provide gene-expression predictions, while CLS representations provide cell-level embeddings for spatial- domain identification. The results therefore support the feasibility of maintaining explicit input-gene correspondence through spatial contextualization while also learning a cell-level summary.

The two representation levels support complementary downstream roles. Contextual gene states support gene-expression reconstruction, whereas the CLS representation provides a cell-level readout for spatial-domain identification. The single-slice ablation further indicates that additional supervision of the cell-level representation is associated with improved clustering, with mean ARI increasing as CLS-conditioned reconstruction and contrastive supervision are added. Post-hoc spatial refinement provides a complementary downstream effect on the Visium datasets where it is applied, indicating that the final domain assignments can additionally benefit from local spatial continuity beyond clustering the learned CLS representations alone.

An important limitation is computational scaling. Explicit cell–gene states increase computation and memory requirements as gene coverage and neighborhood-expanded context grow, and dense within-cell attention scales quadratically with the number of genes. The current neighborhood-expanded mini-batch scheme also restricts spatial information exchange to the local context included in each batch, limiting recursive propagation of information across multiple graph hops. In addition, the contrastive objective relies on positive and comparison sets constructed within each batch, where local spatial and expression relationships may not always provide unambiguous positive– negative assignments. These limitations motivate future work on more efficient computation, broader spatial-context propagation, and more robust construction of contrastive relationships.

Overall, SpaAxFormer provides a unified gene-resolved formulation for combining within-cell and spatial context while retaining both gene-level and cell-level representations. The study shows that these representation levels can be used within the same framework for gene-expression reconstruction and spatial-domain identification, while the observed behavior highlights the importance of how spatial context and supervision are incorporated.

## Data availability

The original DLPFC data are publicly available through spatialLIBD and the HumanPilot resource at https://github.com/LieberInstitute/HumanPilot [17, 18].

BC1 corresponds to the 10x Genomics Human Breast Cancer (Block A Section 1) dataset [1]. The evaluation annotations used for BC1 are available in *BC*.*zip* distributed by BenchmarkST at *https://doi.org/10.5281/zenodo.10698903* [11]; BenchmarkST credits SEDR as the upstream annotation source [30].

STARMap data are from the mouse visual cortex data set described in the original STARMap study [29]. The processed input used in our experiments, *STAℛmap 20180505 BY3 1k*.*h5ad*, is available in the *dataset/STAℛmap/* directory of the ADEPT repository at *https://github.com/maiziezhoulab/ADEPT* [12].

The imputation experiments follow the Impeller benchmark [9]. The public processed-data distribution for the DLPFC, Stereo-seq, and Slide-seq V2 benchmarks is available at *https://doi.org/10.5281/zenodo.11212604*. The original Stereo-seq and Slide-seq V2 datasets are described in their respective source publications [5, 22].

## Code availability

The source code for SpaAxFormer is publicly available at *https://github.com/uci-cbcl/SpaAxFormer*.

